# Sensory experience-induced plasticity of somatostatin interneurons occurs through two-pore potassium (K_2P_) channels

**DOI:** 10.64898/2026.11.06.764984

**Authors:** Briana M. Bohannon, Nellie L. Alonso, Mackenzie G. Walker, Michael Wang, Katharine Kaimaktchiev, Cheryl Yang, Daniel E. Feldman

## Abstract

GABAergic interneurons (INs) are powerful modulators of plasticity and learning in cortical networks, but plasticity of INs themselves is not well understood. We studied experience-dependent plasticity in somatostatin (SST) INs, which critically regulate dendritic integration, sensory tuning, and plasticity in pyramidal neurons in sensory cortex. Whether SST INs exhibit experience-dependent plasticity, and by what mechanisms, is unknown. We found that 3 days of whisker deprivation strongly downregulated intrinsic excitability of SST INs in L2/3 of somatosensory cortex, by reducing input resistance. Deprivation also increased mEPSCs onto SST INs. The reduction in intrinsic excitability was caused by an increase in leak potassium current through two-pore K^+^ (K_2P_) channels, without changes in HCN, K_V_1, or K_V_7 currents, which contribute to plasticity in other IN types and brain areas. Immunohistochemistry showed an increase in TRESK, but not other K_2P_ channels, indicating a novel role for K_2P_ channels in experience-dependent plasticity of interneuron excitability.

## Introduction

In cerebral cortex, GABAergic interneurons (INs) powerfully shape network activity and neural coding, but their roles in experience-dependent plasticity remain only partially understood. Among the 3 major classes of GABAergic INs - somatostatin (SST), parvalbumin (PV), and 5HT3 receptor (5HT3R)-expressing classes - how SST INs contribute to cortical plasticity is least understood. SST INs are highly abundant, making up 30% of the cortical IN population and about 5% of all cortical neurons^1,2^. SST INs preferentially target the distal dendrites of pyramidal (PYR) cells and modulate dendritic integration, local dendritic spiking, and long-term synaptic plasticity^3,4^. Within the SST IN class there is considerable morphological, biophysical, and transcriptomic diversity. Morphologically, SST cells include Martinotti cells (MCs) which have a prominent axonal arborization in L1, and non-Martinotti cells (non-MCs) that have largely local axonal ramifications. Transcriptomic methods identify 8-9 subtypes of SST INs within L2/3 alone^5,6^. The most abundant transcriptomic SST subtype in L2/3 expresses *Calb2*, which encodes the calcium binding protein calretinin (CR), and corresponds to non-bursting Martinotti SST INs^5^. Other transcriptomic subtypes in L2/3 include *Tac1*-expressing SST cells, which are quasi-fast spiking (FS) or “FS-like” SST INs, and *Chodl*-expressing SST cells which are long-range projecting INs that express neuronal nitric oxide synthase (nNos)^5,7^. How SST cells overall, or any of these specific subtypes, contribute to cortical plasticity remains poorly understood.

In sensory cortex, dynamic modulation of SST circuit function, either developmentally^4^, or acutely by VIP neuron activation^8–10^, regulates plasticity in excitatory circuits. Plasticity also alters some features of SST circuits themselves. For example, behavioral training on motor and sensorimotor learning tasks produces sustained alterations in SST circuit function in L2/3 of motor and sensory cortex, which may be a key step in enabling learning-related plasticity in PYR networks^11^. Similarly, visual experience increases visually-evoked activation of SST cells in V1^12^. These learning-related changes involve plasticity at excitatory synapses onto SST cells, or SST inhibitory output synapses^11,13–15^. Plasticity of intrinsic excitability (IE) is a major cellular plasticity mechanism within other interneuron types^16^, but whether SST INs exhibit plasticity of IE in response to experience is unknown.

We tested whether sensory experience drives plasticity of IE in SST cells, and if so, by what mechanisms. Outside of sensory cortex, hippocampal O-LM neurons, a well-studied SST IN subtype, exhibit robust plasticity of IE, in the form of long-term potentiation (LTP) and long-term depression (LTD) of IE in response to brief stimulated activity patterns in brain slices^17,18^. LTD of IE in O-LM cells is mediated by increased synthesis of the endocannabinoid 2-arachidonoylglycerol (2-AG), which increases K_V_7.2/3 current (I_M_) to reduce cell excitability^17^. LTP of IE in O-LM cells occurs in response to 5 Hz (theta burst) stimulation applied to O-LM afferents^18^ and is mediated by a reduction of K_V_7.2/3 and HCN channel currents (I_M_ and I_h_) that both result from membrane depletion of PIP_2_^18^. For neocortical SST INs, plasticity of IE has been reported in slice culture^19^ in response to blockade of network activity using tetrodotoxin (TTX), which increases SST IE due to reductions in I_h_ and a putative leak K^+^ conductance^19^. But whether neocortical SST INs exhibit plasticity of IE in response to sensory experience in vivo, and whether this utilizes these known mechanisms, or novel cellular mechanisms, remains unknown. Here we studied mechanisms of IE plasticity in SST INs in somatosensory cortex (S1), induced by whisker deprivation, a classical manipulation that induces robust experience-dependent plasticity in S1.

## Materials and Methods

### Animal Care

Procedures were approved by the University of California, Berkeley Animal Care and Use Committee and followed National Institutes of Health guidelines. The following mouse strains were utilized: SST-IRES-Cre (Jax #013044), Cre-dependent tdTomato reporter (Ai14; Jax #007914), and GIN (Jax #003718). Male and female mice were used for all experiments. Mice were weaned at P21 into cages of same-sex littermates.

### Whisker deprivation

For whisker deprivation experiments, mice at postnatal day 17-19 (P17-19) were briefly anesthetized with isoflurane (2% in 1 L/min O2) and the D-row whiskers (D1-5 and gamma) were plucked by gentle tension with forceps under an operating microscope. Sham-deprived littermates underwent only anesthesia, and their whiskers were left intact. The plucking or sham procedure took ∼3 minutes per mouse, and mice recovered from anesthesia after 1-2 minutes and were replaced in the home cage. Naïve mice did not undergo anesthesia and had normal whisker experience. Mice that were spontaneously lacking whiskers were not used for experiments.

### Brain slice preparation

Animals were anesthetized using isoflurane and then rapidly decapitated. For mice aged P15-P29, brains were dissected in ice cold, oxygenated (95% O_2_/5% CO_2_) high sucrose cutting solution (75 mM sucrose, 85 mM NaCl, 25 mM D-glucose, 25 mM NaHCO_3_, 2.5 mM KCl, 1.25 NaH_2_PO_4_-H_2_O, 0.5 mM ascorbic acid, 4 mM MgCl_2_, 0.5 mM CaCl_2_; pH = 7.2; 320 mOsm). For mice >P30, slices were prepared in an NMDG cutting solution (135 mM NMDG, 23 mM NaHCO_3_, 1.5 mM KH_2_PO_4_, 0.4 mM ascorbic acid, 1.5 mM KCl, 10 mM D-glucose, 3.5 mM MgCl_2_, 0.5 mM CaCl_2_; pH = 7.4; 320 mOsm). Brain slices (350 μm thickness) were cut on a vibratome (Leica VT1200S) in a plane of section that allows visualization of the A-E barrels, termed the “across-row” slice plane, angled 45° toward coronal from the mid-sagittal plane^20^. Slices were placed in a resting chamber containing oxygenated (95% O_2_/5% CO_2_) standard Ringer’s solution (119 mM NaCl, 26.2 mM NaHCO_3_, 11 mM D-glucose, 1.3 mM MgSO_4_-7H_2_O, 2.5 mM KCl, 1 mM NaH_2_PO_4_-H_2_O, 2.5 mM CaCl_2_; pH = 7.3, 300 mOsm) and maintained for 30 minutes at 32° C followed by 0.5-6 hours at room temperature prior to electrophysiological recordings.

### Electrophysiological recordings

Recordings were made from either L2/3 SST INs or L2/3 PYR cells. All SST recordings except Fig. 2G-I were made in SST-Cre;Ai14 mice and were targeted to tdTomato-expressing neurons. For Fig. 2G-I, recordings were made in GIN mice and targeted to GFP-expressing neurons. Barrels in layer 4 were visualized by transillumination to identify the D-barrel columns in each slice. All electrophysiological recordings were made in the D-barrel column.

#### Current-clamp recordings of SST IN intrinsic excitability

To measure the intrinsic excitability of SST INs, we performed whole-cell current-clamp recordings from L2/3 SST INs in the D-barrel column. Borosilicate glass pipettes (3-5 MΩ) were filled with potassium-gluconate internal solution (116 mM potassium-D-gluconate, 20 mM HEPES, 6 mM potassium chloride (KCl), 2 mM sodium chloride (NaCl), 0.5 mM EGTA, 4 mM Mg-ATP, 0.3 mM Na-GTP, and 10 mM phosphocreatine, pH = 7.25, 290 mOsm). In current-clamp experiments, bridge balance was applied to compensate for series resistance artifacts. Resting membrane potential (V_rest_) was measured immediately upon break-in and cells were allowed to rest freely at V_rest_ for frequency-current (F-I) curve, rheobase, and input resistance measurements. Depolarizing current injections (square pulse, 500 ms duration) were applied to evoke spikes. Rheobase was defined as the minimum current required to evoke a single spike. Rheobase was determined using an ascending and descending ladder protocol with a resolution of 10 pA. F-I curves were measured using ascending current steps starting at rheobase and ending at 200 pA above rheobase (20 pA increments). Input resistance (R_in_) was measured by using -10 pA, 0 pA, and +10 pA current injections (500 ms duration). To measure voltage-sag (HCN-dependent membrane depolarization), cells were held at -65 mV, from which 500 ms hyperpolarizing square pulses were applied (from -50 pA to -800 pA in 50 pA increments). Voltage sag was measured at current steps where peak hyperpolarization reached -100 mV. Action potential bursts were defined as spike patterns that showed both burst acceleration (decreasing inter-spike interval [ISI] accompanied by decreasing spike amplitude) and post-burst deceleration (increasing inter-spike interval accompanied by increasing spike amplitude). Burst onset was defined by the presence of at least 3 accelerating spikes in which the 2nd ISI was at least 3 ms shorter than the first ISI, accompanied by a spike amplitude decrease of at least 1 mV. Burst offset was defined by a corresponding pattern of spike deceleration. Burst index was defined as the sum, across all spikes within a burst, of sqrt(ΔSpikeHeight^2+ΔISI^2), where ΔSpikeHeight is the change in spike amplitude (mV) between this spike and the preceding spike, and ΔISI is the change in ISI (ms) between this ISI and the preceding ISI. This metric is zero in non-bursting cells and increases with the magnitude of acceleration and deceleration within a burst, the number of spikes in the burst, and the number of bursts within the sweep.

#### Voltage-clamp recordings of miniature excitatory and inhibitory post-synaptic currents (mEPSCs/mIPSCs)

To measure mEPSCs and mIPSCs in L2/3 SST INs, we performed whole-cell voltage-clamp recordings using a cesium-gluconate internal solution (108 mM D-gluconic acid, 108 mM cesium hydroxide (CsOH), 20 mM HEPES, 5 mM tetraethylammonium chloride (TEA-Cl, 2.8 mM NaCl, 0.4 mM EGTA, 4 mM Mg-ATP, 0.3 mM Na-GTP, 10 mM phosphocreatine, and 5 mM QX314 bromide, pH = 7.20, 290 mOsm). Series resistance (R_s_) was measured throughout the voltage-clamp recording and were compensated up to 30%. Cells with R_s_ > 20 MΩ were excluded from analysis. mEPSCs and mIPSCs were isolated by holding cells at -70 mV and 0 mV, respectively, each holding potential corrected for junction potential (12 mV). Miniature events were collected in 10 sec long sweeps and data was sampled at 10 kHz and filtered at 3 kHz.

#### Voltage-clamp recordings of K_V_7.2/7.3 channel currents (I_M_)

For voltage-clamp recordings of I_M_, Ringer’s solution contained 400 nM TTX, 10 μM NBQX, 50 μM D-AP5, 3 μM gabazine, and 1 μM ZD7288. Whole-cell recordings were made using potassium-gluconate internal containing 5 mM QX-314, 4 Mg-ATP, 0.3 Na-GTP, and 10 phosphocreatine. In M-current recordings, the addition of Mg-ATP, Na-GTP, and phosphocreatine is crucial for the prevention of membrane PIP_2_ depletion and subsequent M-current run-down^21,22^. To isolate neuronal M-current, cells were held at - 60 mV and stepped to -15 mV for 500 ms to activate outward K^+^ current before subsequent tail currents were measured at -40 mV (1 sec). Cells were then stepped back to -15 mV for 500 ms before returning to -60 mV^17,18^.

#### Voltage-clamp recordings of potassium leak current (I_K(Leak_)

For voltage-clamp recordings of I_K(Leak),_ Ringer’s solution contained 400 nM TTX, 10 μM NBQX, 50 μM D-AP5, 3 μM gabazine, 10 μM XE-991, 100 μM CdCl_2_, 50 μM NiCl_2_, and 1 μM ZD7288. Cells were held at -60 mV in whole-cell voltage-clamp before applying a 6 sec depolarization to -20 mV to establish a standing outward current. Inward K^+^ current was then evoked by applying a series of 3 sec hyperpolarizing test pulses. Current amplitudes were measured at the end of the test pulse to ensure currents have reached steady state. Data was sampled at 10 kHz and filtered at 3 kHz. To isolate the K_2P_-dependent leak K^+^ current, we bath applied 100 μM Ba^2+^ and repeated I-V curve measurements.

### Immunohistochemistry

Animals were anesthetized using isoflurane and underwent transcardial perfusion with ice cold phosphate buffered saline (PBS, Gibco: 10010-023) followed by 4% paraformaldehyde (PFA) in 1X PBS. Following perfusions, brains were rapidly dissected and post-fixed for 2 hours in 4% PFA in PBS before being transferred to 30% sucrose in PBS and stored at 4° C. Histological sections (50 μm thickness) were cut in the same “across row” plane as for slice physiology. Sections were prepared using a sliding microtome and stored in 0.1 M phosphate buffer (PB) prior to immunofluorescence co-labeling experiments. *For AnkG and K_V_1.1 staining:* Sections were washed 3 x 5 minutes in 1X PBS, prior to blocking for 1 hour at room temperature in BlockAid (Invitrogen). The following primary antibodies were used: anti-calretinin (1:500, Millipore-Sigma: MAB1568), anti-K_V_1.1 (1:500, Millipore-Sigma: MABN616), anti-AnkyrinG (1:500, Millipore-Sigma: MABN466). Primary antibodies were diluted in BlockAid with 0.1% TritonX-100 and applied overnight at room temperature on an orbital shaker. The following secondaries antibodies were used: anti-mouse IgG1 AlexaFluor 647 (1:250, Invitrogen: A-21240), anti-mouse IgG2b AlexaFluor 488 (1:250, Invitrogen: A21242), anti-mouse IgG2a 488 (1:500, Invitrogen: A-21131). Secondary antibodies were diluted in BlockAid with 0.1% TritonX-100 and applied overnight at 4° C protected from light. *For K_2P_ channel staining (anti-TREK-1, anti-TASK-1, anti-THIK-2, and anti-TRESK):* Sections were washed 3 x 5 minutes in 1X PBS, prior to blocking for 1 hour at room temperature in Blocking Solution (10% normal goat serum, 0.3% Triton-X in 1X PBS). The following primary antibodies were used: mouse IgG1 anti-calretinin (1:500, Millipore-Sigma: MAB1568), guinea pig anti-TREK-1 (1:400, Alomone Labs: APC-047-GP), mouse IgG2b anti-TASK-1 (1:500, Antibodies Inc: 75-357), rabbit anti-THIK-2 (1:400, Alomone Labs: APC-169, and mouse IgG2b anti-TRESK (1:500, Antibodies Incorporated: 75-475). Primary antibodies were diluted in Blocking Solution and applied overnight at room temperature on an orbital shaker. The following secondary antibodies were used: anti-mouse IgG1 AlexaFluor 488 (1:500, Invitrogen: A-21121), anti-guinea pig AlexaFluor 647 (1:250, Invitrogen: A-21450), anti-rabbit AlexaFluor 633 (1:250, Invitrogen: A-21070), anti-mouse IgG2b AlexaFluor 647 (1:250, Invitrogen: A-21242). Secondary antibodies were diluted in Blocking Solution and applied for 2 hours at room temperature, protected from light. Sections were counterstained with 500 nM 4’,6-diamidino-2-phenylindole (DAPI, Invitrogen: D1306) for 20 minutes at room temperature. Sections were then mounted using Prolong Gold Antifade Mounting Media (Invitrogen: P36930). Sham and deprived tissue sections were run in parallel during IHC experiments to control for batch effects so that, within each batch of staining, fluorescence intensity was normalized to levels in sham tissue of the same run.

### HCR RNA-FISH

Animals were anesthetized using isoflurane and underwent transcardial perfusion with ice cold phosphate buffered saline (PBS, Gibco: 10010-023) followed by 4% paraformaldehyde (PFA) in 1X PBS. Following perfusions, brains were rapidly dissected and post-fixed for 24 hours in 4% PFA in PBS before being transferred to 30% sucrose in PBS and stored at 4° C. Histological sections (100 μm thickness) were cut in the same “across row” plane as for slice physiology. Sections were prepared using a sliding microtome and stored in 0.1 M phosphate buffer (PB) prior to immunofluorescence co-labeling experiments. Sections were incubated in 70% EtOH for 2-7 days at 4° C prior to tissue clearing. Clearing was accomplished by incubating tissue sections for 2-3 hours at room temperature in 8% SDS. HCR RNA-FISH reagents from Molecular Instruments were used for HCR-FISH experiments. Floating sections were pre-hybridized in 200 uL of HCR Probe Hybridization Buffer for 30 minutes at 37° C prior to overnight hybridization with probes diluted in HCR Probe Hybridization Buffer at 37° C. The following Molecular Instruments probes were used: *Sst (3 nM), Pvalb (16 nM), Calb2 (16 nM), Kcnk2 (16 nM), Kcnk3 (16 nM), Kcnk12 (16 nM)*, and *Kcnk18 (16 nM)*. Following hybridization, sections were washed 45 min each in pre-warmed 100% HCR Probe Wash Buffer, 75% HCR Probe Wash Buffer/25% 2X sodium chloride sodium citrate with 10% Tween (SSC-T), then 50% HCR Probe Wash Buffer/50% 2X SSC-T. Sections were pre-amplified for 30 minutes at room temperature in HCR Amplification Buffer. Fluorescent hairpins (594, 488, and 647) were snap-cooled for 90 seconds at 95 C. Just prior to amplification, hairpins (at 150 nM) were diluted in HCR Amplification Buffer. Sections were incubated in HCR Amplification Buffer with added hairpins overnight at room temperature, protected from light. Prior to mounting, sections were washed 3 x 1 hr in 2X SSC-T, followed by 3 x 10 min at 2X SSC. DAPI was added in the final SSC washes in the first round for visualization of whisker barrel columns. Sections were then mounted using Prolong Gold Antifade Mounting Media (Invitrogen: P36930). Sham and deprived tissue sections were run in parallel during HCR-FISH experiments to control for batch effects so that, within each batch of staining, fluorescence intensity was normalized to levels in sham tissue of the same run.

### Image acquisition and analysis

Confocal z-stacks with tile scanning were collected on a laser scanning confocal microscope (Zeiss LSM-780). Image post-processing and fluorescence intensity quantification were performed using FIJI (ImageJ) software. Fluorescence intensity was measured in SST INs of L2/3 within the D-barrel column. The cells analyzed were divided into different experimental cell classes: SST cells were all cells that expressed genetically encoded tdTomato; CR+ cells co-expressed SST and CR; CR-cells expressed SST but did not co-express CR. Cellular regions of interest (ROIs) for each cell group were drawn manually. Fluorescence intensity was measured in each of the three cell classes in both sham and D-row deprived (as well as sham and B-row spared) experimental groups and averaged by histological section. For each cell class, the fluorescence intensity of the ROIs in each section were averaged to obtain an average fluorescence intensity per histological section. For normalization, we calculated the average fluorescence intensity per round of IHC by averaging the fluorescence intensities of each tissue section belonging to the same batch. To evaluate relative changes in fluorescence intensity across experiment groups, each cell class in sham vs deprived mice were divided by this average per round from sham SST INs. With this analysis, the n represents number of tissue sections.

### Data analysis and Statistics

Data analysis was performed using Matlab. All statistical comparisons were computed using GraphPad Prism (GraphPad Software, La Jolla, CA). The significance threshold was α=0.05.

## Results

### Developmental maturation of intrinsic excitability in L2/3 SST INs

We first assessed the normal maturation of electrophysiological properties of L2/3 SST INs in L2/3 of S1^4,9,23^. To do this, we made fluorescence-targeted whole-cell current-clamp recordings from SST INs in slices from SST-Cre;Ai14 mice, in which SST cells express tdTomato. Experiments were performed between P15-P60 (Fig. 1A), with synaptic blockers present. SST INs display extensive electrophysiological diversity both within and between transcriptomic and morphological subtypes^5^. Consistent with this, we observed considerable diversity in current-evoked spike patterns, including accommodating (ACC), burst accommodating (bACC), irregularly bursting (IB), and quasi-fast spiking (FS-like) firing patterns, as previously reported for SST INs^24–27^ (Fig 1B).

**Figure 1:**
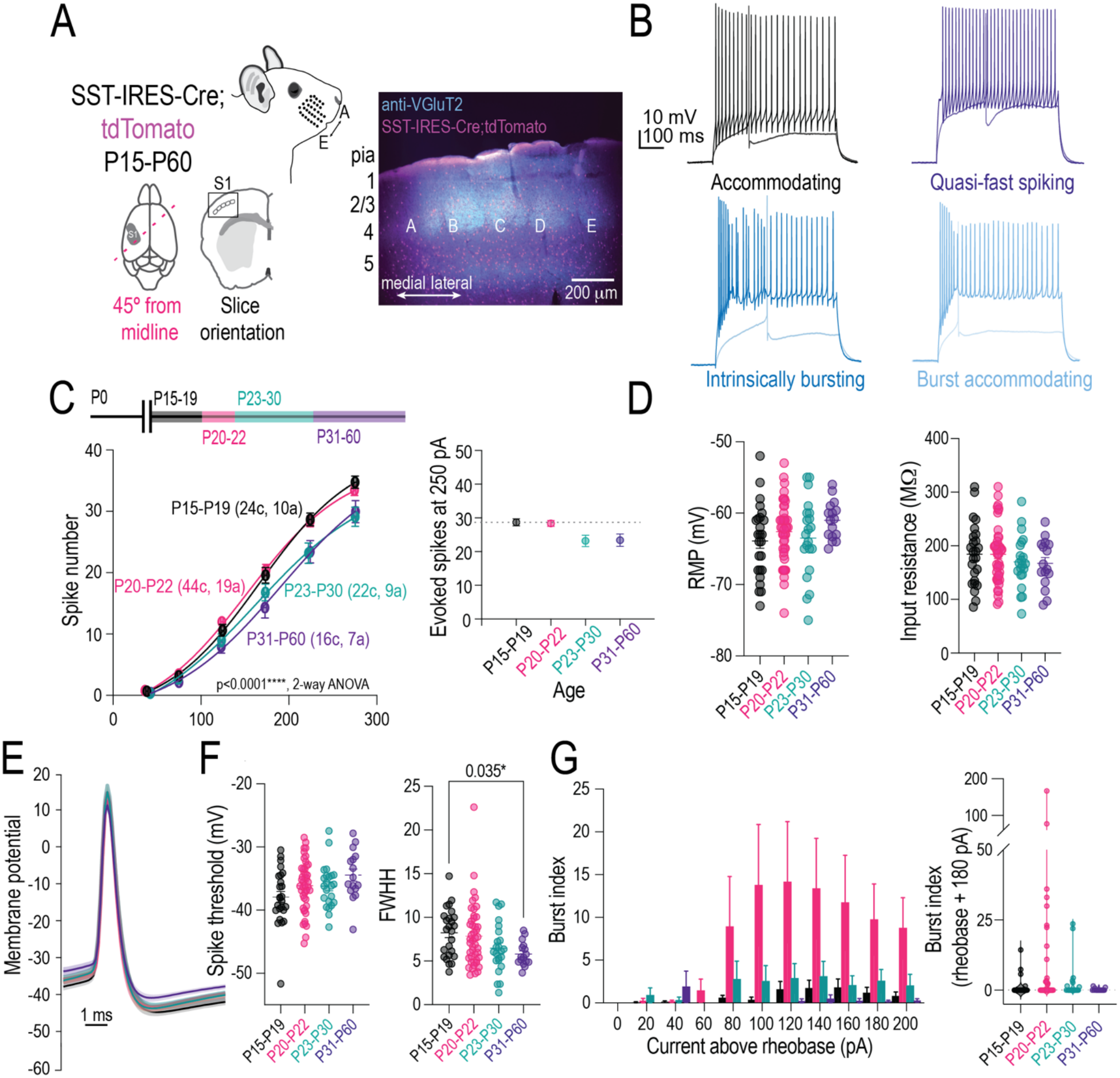
Developmental maturation of intrinsic excitability in L2/3 SST INs. **A.** Schematic of mouse facial whiskers, across-row slice preparation, and organization of whisker barrel columns in SST-IRES-Cre; tdTomato mice. **B.** Representative traces of electrophysiological diversity of L2/3 SST INs. **C.** Spike frequency-current (F-I) relationship (left) and average spike waveform (right) comparing different developmental time bins P15-P19 (black), P20-P22 (pink), P23-P30 (teal), and P31-P60 (dark purple). F-I curves were compared using 2-way ANOVA, p < 0.0001 ****. **D.** Comparison of resting membrane potential (V_rest_) across different developmental time bins (1-way ANOVA, p=0.274, ns) and input resistance (1-way ANOVA, p>0.999, ns). **E.** Average spike waveforms across development. **F.** Comparison of spike threshold) across different developmental time bins (1-way ANOVA, p=0.068, ns) and full width at half height (1-way ANOVA, p = 0.035 *). **G.** Comparison of burst index across different developmental time bins and increasing current injection amplitude (left) as well as comparison of burst index at 180 pA above rheobase (right, 1-way ANOVA, p = 0.098, ns).

For each neuron, we determined rheobase and measured spike frequency-current (F-I) curves for at multiple current steps relative to rheobase. F-I curves showed higher evoked firing rate at young ages (P15-19 and P20-22), that shifted to lower firing rates as animals mature (P23-30 and P31-60) (Fig. 1C). Thus, mean SST IN intrinsic excitability decreased with development. In contrast, the mean spike waveform was relatively constant across development, with the exception of an age-dependent narrowing of the action potential full width at half height (FWHH - P15-P19: 0.82 ± 0.55; P20-P22: 0.79 ± 0.53; P23-P30: 0.64 ± 0.59; P31-P60: 0.58 ± 0.34; Kruskal Wallis test: p = 0.026*, Fig 1E). Most other cell-intrinsic properties, such as resting membrane potential (V_rest_), input resistance (R_in_), rheobase, and spike threshold, were stable across these ages, although R_in_ and spike threshold showed non-significant trends in the direction of reduced cellular excitability with age (Fig. 1D-E, Table 1;).

**Table 1.** Cell-intrinsic properties of L2/3 SST cells (mean ± SEM)

| **Age** | **V<sub>rest</sub> (mV)** | **Input resistance (**MΩ) | **Rheobase (pA)** | **Spike threshold (mV)** | **N(cells)** |
|---|---|---|---|---|---|
| P15-19 | -63.88 ± 1.03 | 184.5 ± 11.60 | 92.75 ± 12.27 | -37.9 ± 0.89 | 25 |
| P20-22 | -62.59 ± 0.63 | 184.1 ± 7.94 | 81.86 ± 7.71 | -36.1 ± 0.60 | 45 |
| P23-30 | -63.48 ± 1.18 | 170.1 ± 10.73 | 91.61 ± 10.20 | -36.2 ± 0.72 | 23 |
| P31-60 | -61.01 ± 0.66 | 167.0 ± 11.16 | 104.3 ± 13.32 | -34.5 ± 0.97 | 16 |

About 30% of L2/3 SST cells responded to current injections with action potential bursts. We quantified bursting using a burst index that reflects patterns of acceleration-deceleration in inter-spike intervals accompanied by variation in spike amplitude. This showed little bursting at P15-19, the appearance of more prominent bursting at P20-22, and then a decline at P31-P60, though differences in burst index across age groups were only trends and not significant (Fig. 1F, P15-P19: 1.159 ± 0.662; P20-P22: 9.749 ± 4.160; P23-P30: 2.800 ± 1.479; P31-P60 0.2031 ± 0.114; Kruskal Wallis test: p = 0.098, ns). Together, these results indicate that SST IN intrinsic excitability is greatest at young ages and declines after the third postnatal week (∼P23), consistent with developmental changes in F-I curves and narrowing of action potential width that have been observed in other cortical cell types^28,29^.

### Whisker deprivation drives reduced intrinsic excitability in L2/3 SST INs

Plasticity of intrinsic excitability (IE) occurs in PYR cells and PV INs in sensory cortex in response to sensory deprivation, and is an important mechanism for homeostatic stabilization of network firing rate^16,30^. In S1, whisker deprivation drives rapid weakening of IE in L2/3 PV cells (which occurs through an increase in expression of the voltage-gated K^+^ channel K_V_1.1 that elevates spike threshold^16,31^) accompanied by a slower weakening of excitatory inputs onto these cells^20^. Altogether, these processes act to reduce inhibition onto PYR cells and homeostatically stabilize L2/3 PYR cell firing rates^32^. Whether SST INs undergo similar experience-dependent plasticity is unknown. To test for experience-dependent plasticity of SST IE, we performed D-row whisker deprivation in SST-IRES-Cre;Ai14 mice beginning at P17-19 and lasting 1-3 days. Following deprivation, we made current clamp recordings in *ex vivo* S1 slices from L2/3 SST INs in D-row whisker columns (i.e., deprived columns). We compared SST IE between these mice and sham-deprived littermates with normal whisker experience (Fig 2A).

**Figure 2:**
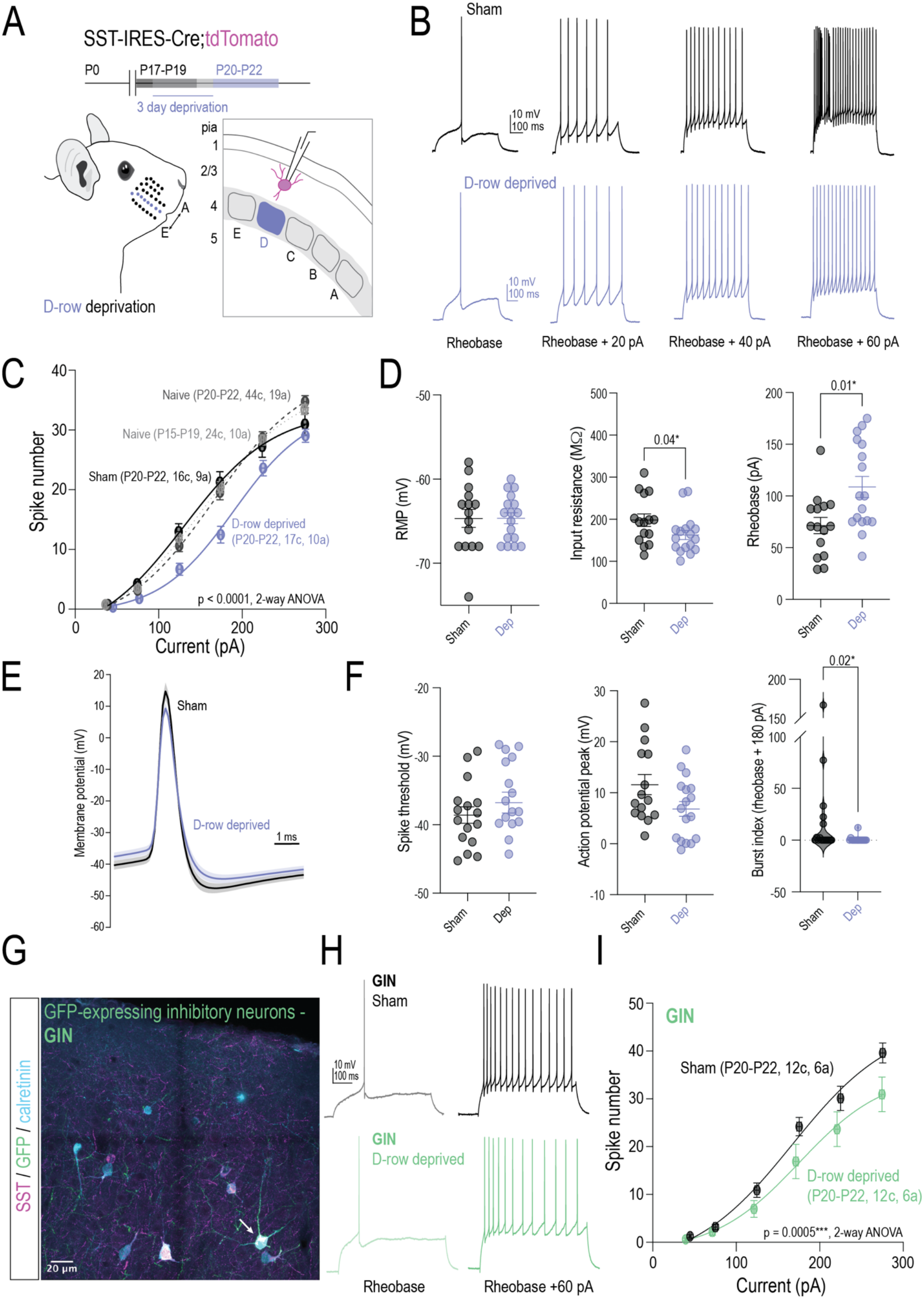
Whisker deprivation drives reduced intrinsic excitability in L2/3 SST INs. **A.** Schematic and timeline of D-row whisker deprivation and targeting of L2/3 SST INs in deprived D barrel columns**. B.** Representative traces of sham and D-row deprived SST INs at rheobase, +20 pA, +40 pA, and +60 pA above rheobase. **C.** Spike frequency-current (F-I) relationship comparing sham (black) and D-row deprived (light purple) mice after D-row whisker deprivation. F-I curves were compared using 2-way ANOVA, p < 0.0001****. **D.** Comparison of cell intrinsic properties including resting membrane potential (V_rest_), input resistance, and rheobase in sham vs D-row deprived mice (Mann-Whitney rank comparisons, V_rest_ p = 0.933, ns, input resistance p = 0.040*, rheobase p = 0.011*). **E.** Average spike waveform in sham vs D-row deprived mice. **F.** Comparison of spiking properties including spike threshold, action potential peak, and burst index (Mann-Whitney rank comparisons, spike threshold p = 0.350, ns, action potential peak p = 0.097, ns, burst index p = 0.021*). **G.** Representative fluorescence co-staining image from GFP-expressing inhibitory neurons (GIN) mice illustrating SST, GFP, and calretinin (CR) expression. **H.** Representative traces from sham and D-row deprived GIN mice. **I.** Spike frequency-current (F-I) relationship comparing sham (black) and D-row deprived (light green) mice after D-row whisker deprivation. F-I curves were compared using 2-way ANOVA, p = 0.005***.

Current-evoked spiking was substantially reduced in L2/3 SST INs in mice after 3 days of D-row whisker deprivation, relative to sham mice (Fig. 2B-C). No reduction in IE was observed after 1 or 2 days of deprivation (Supp. Fig. 1). The reduction in IE after 3 days of deprivation was associated with reduced input resistance (deprived: 162.7 ± 11.0 MΩ, n=17 cells, 10 mice, sham: 197.9 ± 14.7 MΩ, n=16 cells, 7 mice, Mann-Whitney test: p = 0.040) and also with increased rheobase (deprived: 108.7 ± 10.2 pA, sham: 71.40 ± 7.8 pA, Mann-Whitney: p = 0.011). In contrast, V_rest_ was unchanged (deprived: -64.65 ± 0.66 mV, sham: -64.67 ± 1.07 mV, Mann-Whitney: p = 0.933) (Fig. 2D). Spike properties such as action potential peak (deprived: 6.82 ± 1.5 mV, sham: 11.6 ± 2.0 mV, Mann-Whitney: p = 0.097) and spike threshold (deprived: -35.75 ± 1.2 mV, sham: -37.41 ± 1.2 mV, Mann-Whitney: p = 0.350) (Fig. 2E-F) were unchanged after whisker deprivation. However, we found significant reductions in action potential bursting after whisker deprivation (Fig. 2F) (burst index for deprived: 0.845 ± 0.71, sham: 20.00 ± 11.0; Mann-Whitney: p = 0.021).

Weaker F-I curves observed after 3 days of deprivation represent an active weakening of IE in response to deprivation, not a failure of normal maturation, because F-I curves in 3-day deprived cells (measured at P20-22) were not only weaker than sham-deprived mice at P20-22, but also weaker than naive mice with normal whisker experience at P15-19 (Fig. 2C) (spike number for deprived: P20-P22: 12.47 ± 1.4; P15-P19: 19.6 ± 1.3; p < 0.0001****). In addition, F-I curves were on average identical between naive mice at P15-19 and P20-22, indicating that F-I curves are not normally strengthening during this developmental window (Fig. 1C).

Therefore, we observe a pattern of unchanged V_rest_, reduced input resistance, increased rheobase, and no change in spike threshold following whisker deprivation. This suggests that reduced excitability primarily reflects the deprivation-induced reduction in input resistance. This differs from deprivation-induced weakening of IE in PV cells, which is mediated by increased spike threshold without changes in input resistance^16^, and suggests a fundamentally distinct biophysical mechanism of plasticity in SST INs compared to PV INs in S1.

It is unclear if the deprivation-induced reduction in SST IE occurs ubiquitously across all SST IN subtypes or in a subtype-specific manner. Therefore, we tested whether deprivation-induced plasticity of IE was present in one subset of L2/3 SST INs by utilizing the GIN mouse line, where GFP is expressed in Martinotti-type SST INs^33,34^ (Fig. 2G). GIN mice were subject to 3-day, D-row whisker deprivation or sham-deprivation at P17-19 as described for the SST-IRES-Cre;Ai14 mice above. We patched GFP+ cells (“GIN cells”) in L2/3 of D columns and assessed F-I curves and passive properties (Fig. 2G-I). GIN cells from L2/3 of sham mice did not show spike bursting, consistent with previous reports^33,35^. This indicates that the bursting SST cells we observed in SST-Cre;Ai14 mice are likely non-Martinotti cells. 3-day deprivation significantly reduced F-I curves in GIN cells in deprived mice relative to sham mice (Fig. 2I). Thus, deprivation reduces IE in Martinotti-type SST INs in GIN mice. Because the magnitude of this effect in GIN mice is similar to the broad population of SST INs in SST-IRES-Cre;Ai14 mice, and because in SST-IRES-Cre mice, deprivation strongly reduces bursting (which is a property of non-Martinotti cells), we conclude that both Martinotti and non-Martinotti SST subtypes show deprivation-induced reduction of IE.

### Whisker deprivation increases mEPSC frequency onto SST INs

We tested whether SST INs exhibit deprivation-induced changes in synaptic inputs, as occurs in PV cells^20,36^ and PYR cells during sensory manipulation-induced homeostatic synaptic scaling^37,38^. We subjected SST-Cre;Ai14 mice to 3 days of D-row whisker deprivation or sham deprivation as above. Following deprivation, we measured synaptic inputs onto L2/3 SST INs by recording miniature excitatory and inhibitory post-synaptic currents (mEPSCs and mIPSCs), separated by holding cells at -70 mV and 0 mV, respectively. For mEPSCs, we observed a non-significant trend for increased mean mEPSC amplitude in deprived mice relative to sham mice (deprived: -18.13 ± 1.7 pA, n = 14 cells, 6 mice, sham: -13.81 ± 0.759 pA, n = 13 cells, 8 mice, Mann-Whitney test, p = 0.056), and a significant reduction in mEPSC inter-event interval (i.e., increased mEPSC frequency) in deprived mice (deprived: 0.358 ± 0.034 sec, sham: 0.613 ± 0.081 sec, p = 0.0067) (Fig. 3A-B). For mIPSCs, no significant changes were apparent in mIPSC amplitude (deprived: 12.24 ± 0.969 pA, sham: 10.57 ± 0.573 pA, p = 0.094) or inter-event interval (deprived: 0.329 ± 0.085 sec, sham: 0.432 ± 0.091 sec, p = 0.116) (Fig. 3C-D). Thus, 3-day deprivation elevated the rate of mEPSCs, suggesting a greater number or higher release probability excitatory synapses onto L2/3 SST INs, but no changes in inhibitory synaptic inputs.

**Figure 3:**
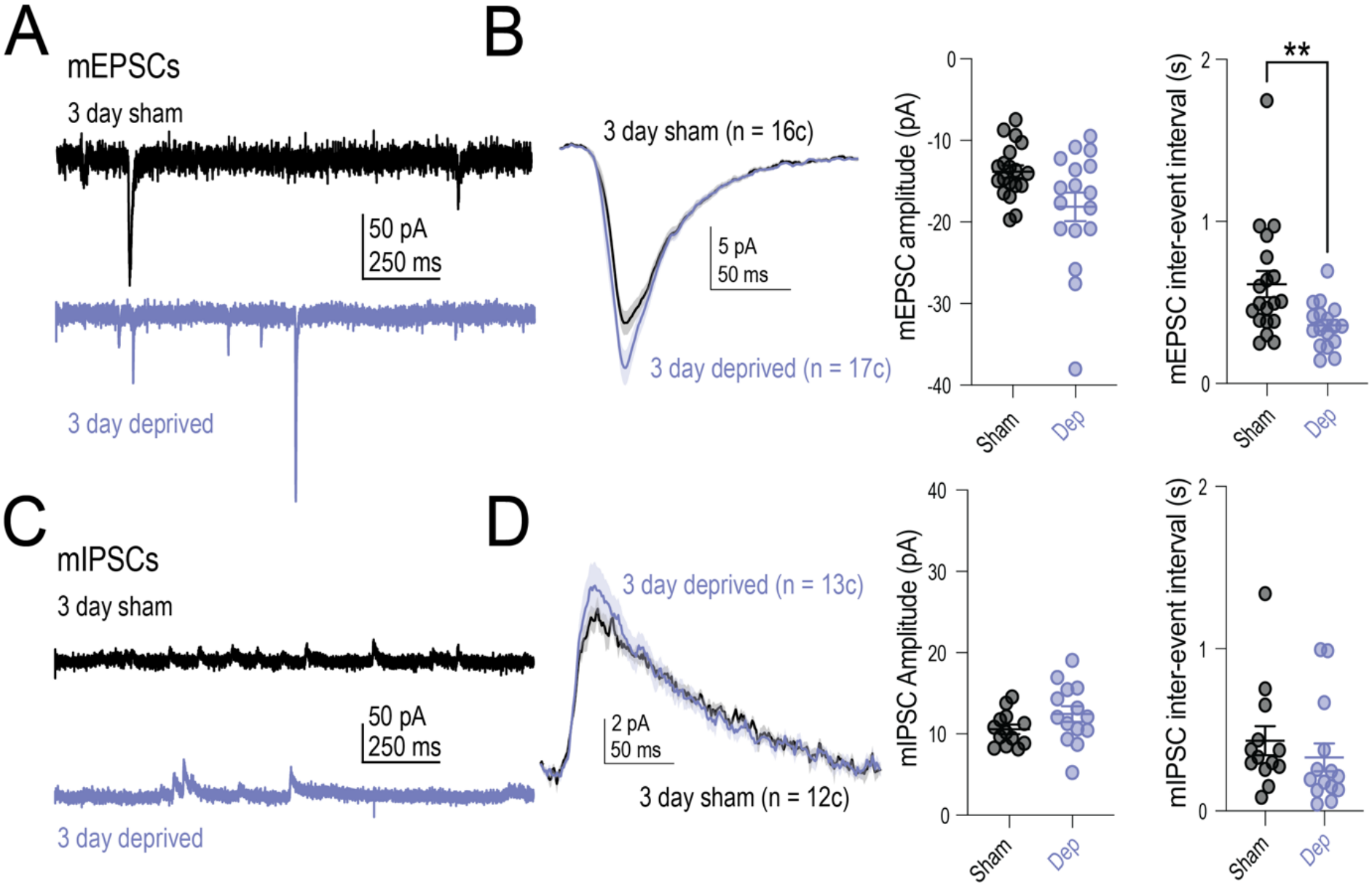
Plasticity of mEPSCs onto SST INs. **A.** Representative EPSCs in sham and D-row deprived mice. **B.** Population average mEPSCs in sham vs D-row deprived mice (left) and quantification of mEPSC amplitude and inter-event interval (IEI, right). **C.** Representative mIPSCs in sham and D-row deprived mice. **D.** Population average mIPSCs (left) and quantification of mIPSC amplitude and inter-event interval (IEI, right).

### Whisker deprivation does not upregulate K_V_1.1 channel expression in SST Ins

Plasticity of IE is typically mediated by altered expression and/or function of ion channel proteins, but the specific molecular and biophysical mechanisms for IE plasticity are diverse and cell-type specific. We sought to elucidate the biophysical mechanisms that mediate the deprivation-induced reduction of SST IE. In PV cells in S1, whisker deprivation reduces PV IE by increasing expression of the voltage-gated K^+^ channel K_V_1.1, detectable at the mRNA and protein level^31^. This increase in channel expression elevates K_V_1 current^16^ and increases spike threshold^16,31^. To test whether SST IE plasticity has this same mechanism, we used IHC to evaluate changes in the expression of K_V_1.1 protein in S1 tissue sections from SST-IRES-Cre;Ai14 mice, where all tdTomato+ cells are SST INs (Supp. Fig. 2A). We co-stained for K_V_1.1 and calretinin (CR), which mark the SST Martinotti cell (*Sst-Calb2*) subtype. tdTomato-positive cells that co-expressed CR (SST/CR+) were classified as Martinotti-like cells, whereas tdTomato-positive cells that did not express CR (SST/CR-) were defined as non-Martinotti-like cells. Fluorescence intensities from each subtype of SST IN (SST, SST/CR+, and SST/CR-) were normalized to the fluorescence intensity of sham SST INs. In sham mice with normal whisker experience, K_V_1.1 was enriched in the non-Martinotti cell population compared to the average SST or Martinotti cell populations (Supp. Fig. 2B, normalized mean K_V_1.1 fluorescence intensity: SST: 1.00 ± 0.04, SST/CR+: 0.951 ± 0.04, SST/CR-: 1.123 ± 0.05, p =0.049*, Kruskal-Wallis test). However, 3 days of whisker deprivation did not alter K_V_1.1 expression in either all L2/3 SST INs, L2/3 SST/CR+ INs, or L2/3 SST/CR-INs (Supp. Fig. 2C, normalized mean fluorescence intensity – SST sham: 0.98 ± 0.03 vs SST deprived: 1.06 ± 0.12; SST/CR+ sham: 0.92 ± 0.03 vs SST/CR+ deprived: 0.97 ± 0.12; SST/CR-sham: 1.09 ± 0.04, SST/CR-deprived: 1.08 ± 0.14). This finding, together with the lack of regulation of spike threshold by deprivation (Fig. 2), suggest that SST INs regulate their intrinsic excitability by a K_V_1.1-independent mechanism.

### Axon initial segment (AIS) structure is unchanged following whisker deprivation

The axon initial segment, which is enriched for the scaffolding protein Ankyrin G (AnkG), is the site of action potential initiation and expresses in Na_V_ and K_V_ channels at high density^39–41^. In PYR cells, altered sensory experience leads to changes in AIS length^30^ that correlate with PYR intrinsic excitability. To test whether SST IE plasticity is associated with AIS changes, we assayed for deprivation-induced changes in AIS morphology, including AIS length and distance from the soma. We analyzed AIS’s of SST INs in L2/3 of D columns from 3-day whisker deprived and sham-deprived SST-IRES-Cre;Ai14 mice. SST INs were defined based on tdTomato+ expression, and Martinotti-like (SST/CR+) and non-Martinotti-like (SST/CR-) subclasses were identified based on co-localization for the marker calretinin (CR). The AIS for these cells was identified by the immunolabeling of AnkG. Irrespective of SST cell subtype, there were no changes in the AIS length between deprived and sham mice after 3-day whisker deprivation (Supp. Fig. 2E, SST sham: 16.32 ± 0.54 μm vs SST deprived: 17.34 ± 0.43 μm, p = 0.14, ns; SST/CR+ sham: 16.17 ± 0.62 μm vs SST/CR+ deprived: 16.68 ± 0.65 μm, p = 0.57, ns; SST/CR-sham: 16.52 ± 1.0 μm vs SST/CR-deprived: 17.76 ± 0.58 μm, p = 0.26, ns). In addition, there were no differences between deprived and sham mice in the distance of the AIS from the soma (Supp. Fig. 2F, SST sham: 7.76 ± 0.76 μm vs SST deprived: 9.98 ± 0.91 μm, p = 0.10, ns; SST/CR+ sham: 8.42 ± 1.15 μm vs SST/CR+ deprived: 11.59 ± 1.56 μm, p = 0.06, ns; SST/CR-sham: 6.84 ± 0.86 μm vs SST/CR-deprived: 8.96 ± 1.09 μm, p = 0.49, ns). Thus, SST IE plasticity does not involve detectable morphological changes in the AIS.

### K_V_7.2/K_V_7.3 (M-current) and HCN currents are not altered by whisker deprivation

The voltage-gated K^+^ channels K_V_7.2 and K_V_7.3 underlie the depolarization-activated K^+^ conductance referred to as the neuronal M-current^42^. M-current is active at subthreshold membrane potentials relative to other K_V_ channels, making it a powerful modulator of cellular excitability and timing between action potentials^21,43–46^. In hippocampal SST cells (O-LM cells), theta burst stimulation that drives network plasticity triggers downregulation of M-current, which together with the downregulation of HCN current, causes IE of O-LM cells to increase^18^. This suggests the activity of K_V_7.2/3 and HCN channels can be functionally upregulated by activity. We used a combination of voltage-clamp protocols (adapted from Sammari et al.^18^ and Incontro et al.^17^) and pharmacology to isolate neuronal M-current and test whether it is upregulated after deprivation. Cells were held at -60 mV and stepped to -15 mV to activate outward K^+^ current before subsequent tail currents were measured at -40 mV. We confirmed that the measured current was indeed M-current by blocking it with the by the selective K_V_7.2/3 antagonist XE-991 (Fig. 4A). We then compared M-current amplitude between L2/3 SST INs in the D column of 3-day D-row whisker-deprived mice and sham mice. M-current amplitude was not different between these groups (Deprived: 86.58 ± 12.98 pA, Sham: 81.93 ± 12.21 pA, p = 0.918, Mann-Whitney test) (Fig. 4B). This suggests that modulation of K_V_7 channels does not mediate deprivation-induced plasticity of SST IE in S1 (Fig. 4B).

**Figure 4.**
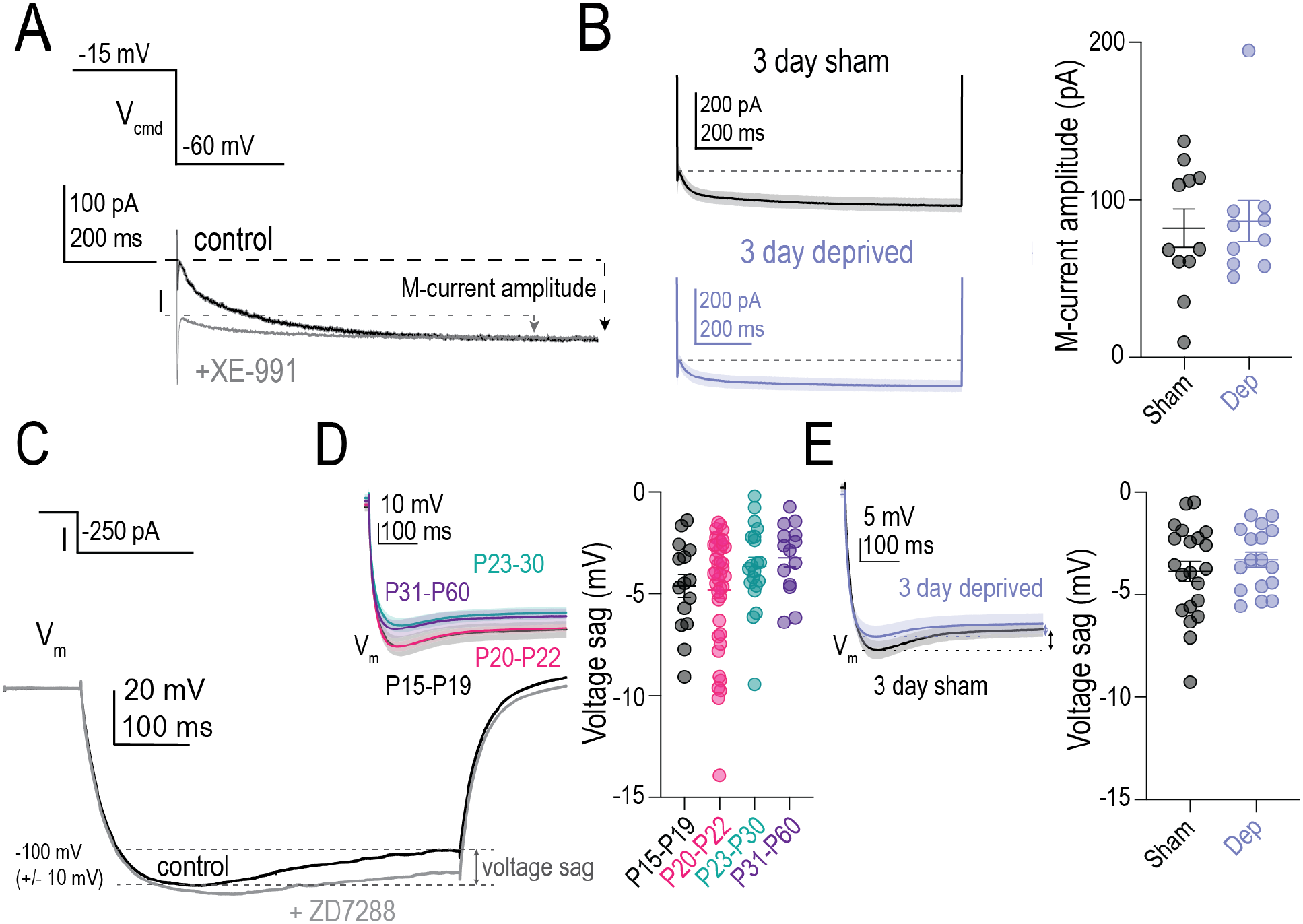
K_V_7.2/K_V_7.3 (M-current) and HCN (H-current) are unaffected by whisker deprivation. **A.** Example trace from individual SST IN of voltage protocol and K_V_7.2/3-dependent M-current (black) and inhibition of M-current by K_V_7.2/3-selective antagonist XE-991 (gray). Dashed lines indicate peak M-current and arrows indicate M-current amplitude. **B.** Comparison of average M-current traces in sham (black, top) vs D-row deprived mice (light purple, bottom) (left) and measured M-current amplitude (right) (Mann-Whitney rank comparison of M-current amplitude p = 0.918, ns). **C.** Example trace from individual SST IN of current protocol and HCN-dependent voltage-sag (black) and inhibition of voltage sag by HCN-selective antagonist ZD7288 (gray). Dashed lines and arrow indicate measured voltage-sag in control trace from peak hyperpolarization to steady state membrane voltage. **D.** Comparison of average voltage-sag traces across different developmental time bins and quantified voltage-sag (P15-P19 - black, P20-P22 - pink, P23-P30 - teal, P31-P60 - dark purple (One-way ANOVA p = 0.16, ns). **E.** Comparison of average voltage-sag traces between sham and D-row deprived mice and quantified voltage-sag (sham - black vs D-row deprived - light purple (Mann-Whitney rank comparison p = 0.399, ns).

We next tested HCN channels, which open at strongly hyperpolarized potentials (-90 to -100 mV), and allow passage of a depolarizing current with slow opening and closing kinetics^47^. HCN channels contribute to resting membrane potential, input resistance, and synaptic integration in many neurons. In O-LM INs, modulation of HCN channels contributes to plasticity of IE following theta burst stimulation^18^. Changes in HCN currents are also implicated in the plasticity of IE in cultured SST INs from S1 at P6^19^. To test whether increased HCN current magnitude contributes to reduced IE and increased input resistance after whisker deprivation in L2/3 SST INs, we performed current clamp recordings and measured voltage sag, which is the characteristic slow depolarization of V_m_ after hyperpolarization onset that is induced by the slow activation of HCN current (Fig. 4C). L2/3 SST INs were held at -65 mV prior to the application of hyperpolarizing current steps. We selected current steps for analysis that yielded a peak V_m_ hyperpolarization to -90 to -100 mV, where HCN channels begin to activate. We confirmed that the measured voltage change was indeed HCN-dependent by blocking it with the selective HCN channel antagonist ZD7288 (1 μM, Fig. 4C). In naive mice with normal whisker experience, voltage sag was present at all ages but showed a trend for lower magnitude in older mice (P15-P19 (black): -4.62 ± 0.57 mV; P20-P22 (pink): 4.80 ± 0.45 mV; P23-P30 (teal): -3.67 ± 0.47 mV; P31-P60 (dark purple): -3.23 ± 0.46 mV; Kruskal-Wallis test p = 0.16, ns; Fig. 4D). We compared voltage sag for L2/3 SST INs at P20-22 in the D-barrel column from 3-day D-row deprived mice with age-matched sham mice. Voltage sag was not significantly different between these groups (Fig. 4E) (deprived (light purple): -3.299 ± 0.371 mV, sham (black): -3.866 ± 0.496 mV; Mann-Whitney test: p = 0.399). Thus, deprivation-induced reduction in SST IE does not appear to be associated with increased HCN current.

### K_2P_-mediated leak potassium current increases with whisker deprivation

Leak potassium current (I_K(Leak)_) is mediated by voltage-independent K^+^ channels and plays a critical role in determining passive membrane properties such as resting membrane potential and input resistance^48^. Whether leak K^+^ channels are modulated by activity or experience to drive plasticity of IE is generally unknown. Changes in I_K(Leak)_ in cortical SST INs have been argued to contribute to IE plasticity in response to chronic activity blockade in slice culture^19^, but whether this occurs under natural conditions *in vivo* is unknown. Because whisker deprivation reduces input resistance in L2/3 SST INs, without changes in HCN current (Fig 4C-E), upregulation of I_K(Leak)_ is a potential mechanism.

To test this, we performed voltage-clamp experiments in L2/3 SST INs using a voltage protocol and pharmacology to isolate I_K(Leak)_. The bath contained synaptic blockers, TTX, XE-991 to block M-currents, ZD-7288 to block HCN currents, and Cd^2+^ and Ni^2+^ to block calcium currents. The holding potential was compensated for junction potential (12 mV). Cells were held at -60 mV before applying a pre-pulse to -20 mV for 6 seconds to allow relaxation of steady-state outward K^+^ current. We then applied 3 second hyperpolarizing voltage steps to measure K^+^ current (Fig. 5A). Steady-state current at the end of the test pulse was linear with V_hold_ between -100 mV and -40 mV, and reversed around -80 mV, consistent with K_leak_ (Fig. 5). At -20 mV, a larger, nonlinear outward current was evident, reflecting the additional contribution of a residual high-threshold voltage-dependent conductance of unknown identity.

**Figure 5.**
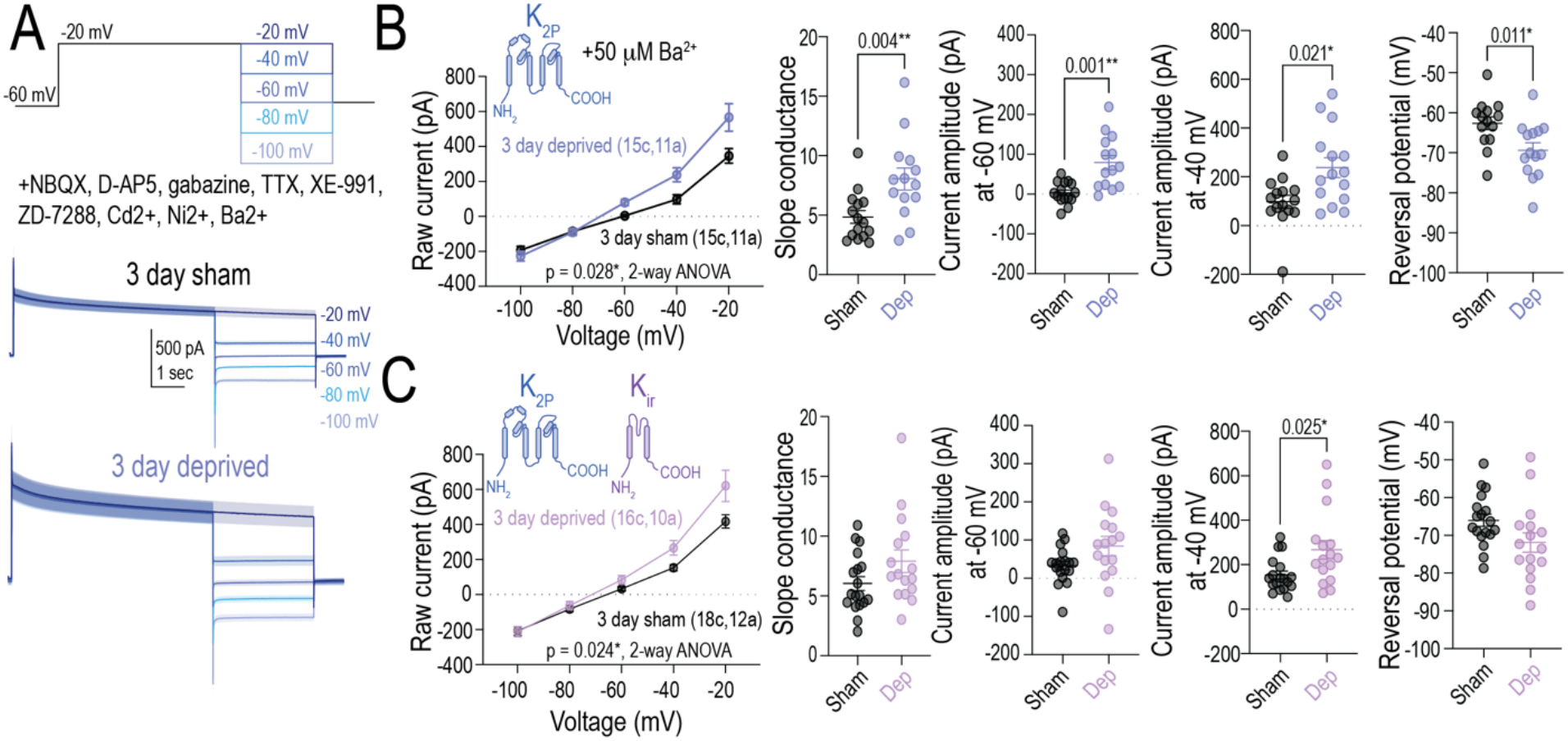
Leak K^+^ current is upregulated in SST INs after whisker deprivation. **A.** Voltage protocol, pharmacology, and population average I_Kleak_ traces in sham and D-row whisker deprived mice. **B.** Current-voltage (I-V) relationship for I_Kleak_ putatively mediated by K_2P_ and K_IR_ channels. Slope conductance measured from fit of the linear component of the I-V curve. Current amplitude at -60 mV holding potential. Reversal potential calculated from linear fit of I-V curve. Explain what p-value represents. **C**. Same as **B,** but for I-V curve measured in 50 μM Ba^2+^ to isolate the K_2P_-mediated component of I_Kleak_.

We tested whether the magnitude of K_leak_ differed between L2/3 SST INs in the D column of 3-day D-row deprived mice vs. sham mice. Two-pore potassium (K_2P_) channels are a major component of outward K^+^ leak current^48^. To isolate the K_2P_ component of leak K^+^ current, we applied a low concentration of Ba^2+^ (which blocks K_IR_ channels, another potential source of K_leak_) and recorded the current vs voltage (I-V) relationship at hyperpolarizing voltage steps (Fig. 5B). We found that this Ba^2+^-insensitive current is significantly upregulated after 3 days of whisker deprivation compared to sham controls (Fig 5B, 2-way ANOVA: p = 0.028*). We also observed a significant increase in leak K^+^ conductance measured by the slope of the I-V relationship (Fig. 5B, D-row deprived: 8.03 ± 0.92 vs sham: 4.84 ± 0.53, Mann-Whitney rank comparison: p = 0.004**). Currents were significantly elevated near resting membrane potential of SST INs (Fig. 5B, -60 mV D-row deprived: 79.1 ± 17.7 pA vs sham: 3.45 ± 7.03 pA, Mann-Whitney rank test: p = 0.001**; and -40 mV D-row deprived: 237.6 ± 40.2 pA vs sham: 97.9 ± 26.5 pA, Mann-Whitney rank comparison: p = 0.02*). In addition, we found a significant hyperpolarization in the reversal potential of measured K^+^ current, suggesting an increase in K^+^ conductance (Fig. 5B, D-row deprived: -69.4 ± 1.86 mV vs sham: -62.6 ± 1.6 mV, Mann-Whitney rank comparison: p = 0.01*).

While K_2P_ channels are the predominant source of outward leak K^+^ current, it is possible that inward rectifier K^+^ channels (K_IR_) can also contribute to K_leak_. The total leak K^+^ current (measured without Ba^2+^ and thus containing K_2P_ + K_IR_) was still significantly elevated following whisker deprivation (Fig. 5C, 2-way ANOVA: p = 0.024*). We observed a strong but non-significant trend towards increased leak K^+^ conductance measured by the slope of the I-V curve (Fig. 5C, D-row deprived: 7.9 ± 0.94 vs sham: 6.03 ± 0.58, Mann-Whitney rank comparison: p = 0.09, ns) as well as K^+^ current measured at -60 mV (Fig. 5C, D-row deprived: 84.4 ± 25.2 pA vs sham: 33.1 ± 11.1 pA, Mann-Whitney rank comparison: p = 0.08, ns). K^+^ current measured at -40 mV was significantly upregulated in D-row deprived compared to sham mice (Fig. 5C, D-row deprived: 266.9 ± 41.9 pA vs sham: 153.2 ± 18.0 pA, Mann-Whitney rank comparison: p = 0.02*). There was also a non-significant trend towards hyperpolarized reversal potential in D-row deprived compared to sham mice (D-row deprived: -71.9 ± 2.6 mV vs sham: -66.0 ± 1.63 mV, Mann-Whitney rank comparison: p = 0.06, ns). Thus, the effect of deprivation is strongest on K_2P_-mediated leak currents, with 3-day deprivation roughly doubling K_2P_-mediated K_leak_ in SST INs, leading to an increase in overall K_leak_ which likely explains the reduction in input resistance and drop in IE.

### Expression of the K_2P_ channel TRESK is upregulated by whisker deprivation

K_2P_ channels are a broad superfamily of K^+^ channels encoded by 15 *Kcnk* genes, subdivided into groups based on structural similarity and gating stimuli (Figure 6A)^49^. The relative expression of these different K_2P_ channel genes in different interneuron types, including SST INs, is not clear. We used immunohistochemistry (IHC) to probe for upregulation of K_2P_ channels in L2/3 SST INs during deprivation-induced plasticity of IE. We focused on candidate K_2P_ channels that we identified in an analysis of publicly available single cell RNA-seq and Patch-Seq datasets, as being expressed in cortical SST INs^50,51^. This analysis revealed that across multiple published datasets, TREK-1, TASK-1, and THIK-2 are enriched in SST INs.

**Figure 6.**
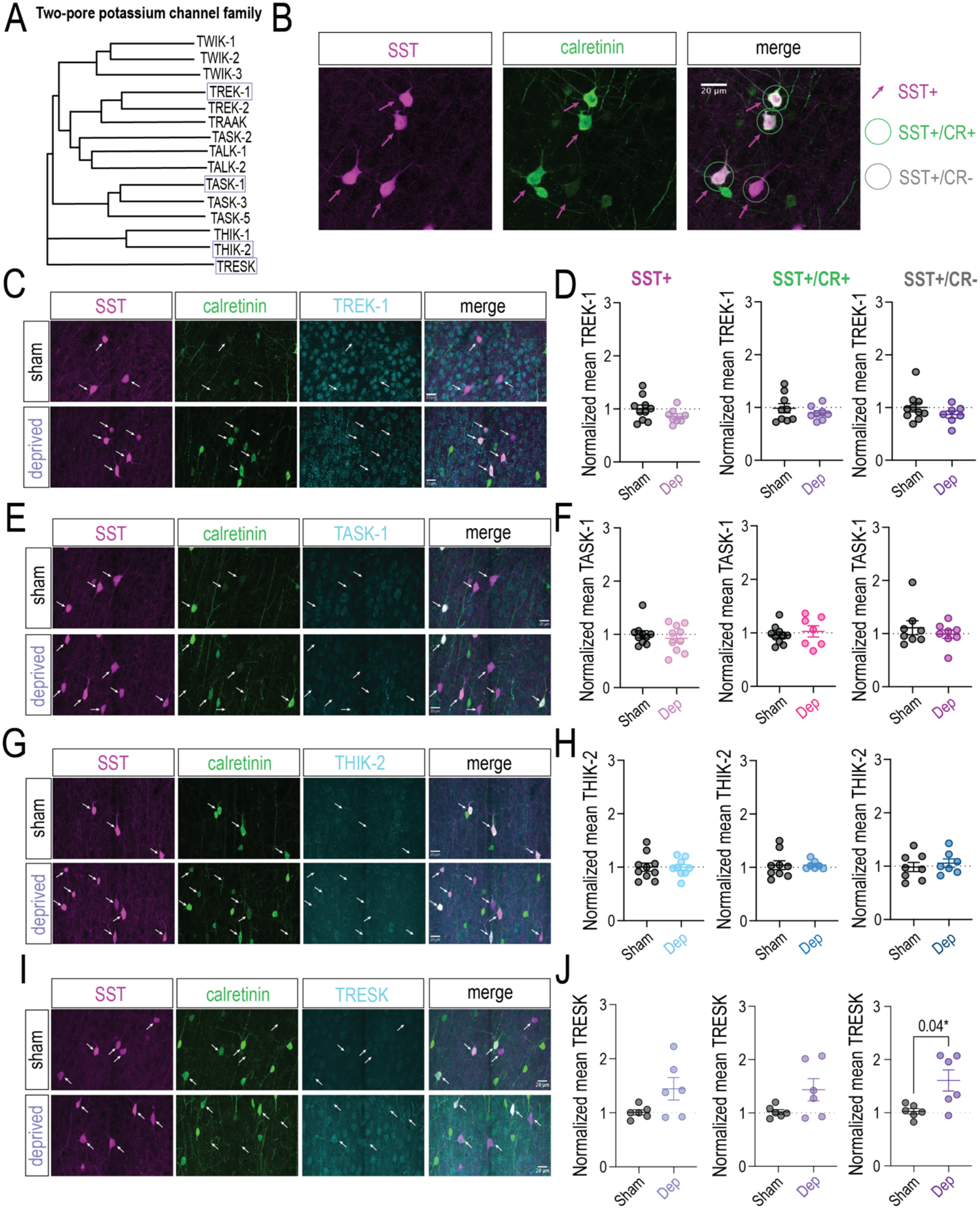
Expression of the two-pore K^+^ channel, TRESK, is upregulated after 3 days of whisker deprivation. **A.** Two-pore K^+^ (K_2P_) channel family, with representative K_2P_ channels whose expression were measured following whisker deprivation highlighted. **B.** Representative image of IHC co-staining for SST and calretinin (CR). Magenta arrows indicate SST INs. Green circles indicate co-localization of SST and CR (putative Martinotti-like SST INs). Gray circles indicate lack of co-localization of SST and CR (putative non-Martinotti-like SST INs). **C.** Representative sham and 3-day deprived co-staining for SST, CR, and the K_2P_ channel TREK-1. **D.** Quantification of mean fluorescence intensity in all SST INs (left), Martinotti-like (SST/CR+) INs (middle), and non-Martinotti-like (SST/CR-) INs (right). **E.** Same as C but for the K_2P_ channel TASK-1. **F.** Same as D but for the K_2P_ channel TASK-1. **G.** Same as C but for the K_2P_ channel THIK-2. **H.** Same as D but for the K_2P_ channel THIK-2. **I.** Same as C but for the K_2P_ channel TRESK. J. Same as D but for the K_2P_ channel TRESK.

To test for regulation of these channels by deprivation, we performed IHC on S1 sections from 3-day deprived and sham SST-IRES-Cre;Ai14 mice, and stained for either TREK-1, TASK-1, or THIK-2, along with CR and SST. SST INs were defined based on tdTomato expression, and Martinotti-like (SST/CR+) and non-Martinotti like (SST/CR-) subclasses were identified based on co-localization for calretinin (CR) (Fig. 6B). TREK-1, which is encoded by *Kcnk2*, belongs to the subclass of K_2P_ channels that are mechanosensitive, temperature sensitive, and activated by polyunsaturated fatty acids and volatile anesthetics^49^. TREK-1 showed the most relative abundance in SST INs compared to other K_2P_ channels^50,51^. When we measured the expression of TREK-1 in sham and deprived mice, we found no deprivation-induced changes in expression in SST INs, Martinotti-like SST INs, or non-Martinotti-like SST INs. (Fig. 6C-D - Sham SST: 1.0 ± 0.07 vs Deprived SST: 0.86 ± 0.05, Mann-Whitney test p = 0.24, ns; Sham SST/CR+: 0.99 ± 0.09 vs Deprived SST/CR+: 0.90 ± 0.05, Mann-Whitney test p = 0.61, ns; Sham SST/CR-: 1.0 ± 0.09 vs Deprived SST/CR-: 0.89 ± 0.07, Mann-Whitney test p = 0.31, ns) TASK-1, encoded by *Kcnk3*, belongs to the acid sensitive subclass of K_2P_ channels^49^. When we measured the expression of TASK-1 in sham and deprived mice, we found no deprivation-induced changes in expression in SST INs, Martinotti-like SST INs, or non-Martinotti-like SST INs. (Fig. 6E-F - Sham SST: 1.0 ± 0.07 vs Deprived SST: 0.93 ± 0.08, Mann-Whitney test p = 0.85, ns; Sham SST/CR+: 0.96 ± 0.06 vs Deprived SST/CR+: 1.03 ± 0.10, Mann-Whitney test p = 0.67, ns; Sham SST/CR-: 1.11 ± 0.13 vs Deprived SST/CR-: 0.99 ± 0.08, Mann-Whitney test p > 0.99, ns.) THIK-2, encoded by *Kcnk12*, belongs to the subclass of K_2P_ channels that are inhibited by halothanes^49^. When we measured the expression of THIK-2 in sham and deprived mice, we found no deprivation-induced changes in expression in SST INs, Martinotti-like SST INs, or non-Martinotti-like SST INs. (Fig. 6G-H - Sham SST: 1.0 ± 0.08 vs Deprived SST: 1.0 ± 0.06, Mann-Whitney test p > 0.99, ns; Sham SST/CR+: 1.04 ± 0.08 vs Deprived SST/CR+: 1.04 ± 0.03, Mann-Whitney test p = 0.68, ns; Sham SST/CR-: 0.98 ± 0.09 vs Deprived SST/CR-: 1.06 ± 0.08, Mann-Whitney test p = 0.54, ns.). Thus, none of the three most commonly expressed K_2P_ genes in SST cells were significantly modulated by deprivation.

We next examined the K_2P_ channel TRESK, encoded by *Kcnk18*. TRESK is a unique member of the K_2P_ family that is activated by rises in intracellular calcium and shares little sequence homology with other K_2P_ genes^52^. TRESK has typically been implicated in pathophysiology leading to pain and migraine^53^, however it has also been demonstrated to play a role in homeostatic plasticity in hippocampal pyramidal cells following seizure-like depolarization events^54^. TRESK is expressed at low levels in cortical SST cells in the publicly available scRNA and Patch-Seq datasets, but these are from animals with normal sensory experience. To test if TRESK may be upregulated after deprivation, we performed IHC on S1 sections from 3-day deprived and sham mice and stained for TRESK and CR while using tdTomato expression to identify SST INs. We found significant elevation of TRESK in non-Martinotti-like (SST/CR-) SST INs after whisker deprivation (Fig. 6I-J - Sham SST/CR-: 0.96 ± 0.13 vs Deprived SST/CR-: 1.51 ± 0.18, Mann-Whitney test p = 0.032*), and a strong trend toward increased TRESK expression in both the full SST IN population and Martinotti-like (SST/CR+) SST INs (Fig. 6I-J - Sham SST: 1.0 ± 0.09 vs Deprived SST: 1.4 ± 0.16, Mann-Whitney test p = 0.16, ns; Sham SST/CR+: 1.01 ± 0.08 vs Deprived SST/CR+: 1.33 ± 0.16, Mann-Whitney test p = 0.26, ns). Thus, the TRESK channel is upregulated by whisker deprivation at the protein level and may contribute to elevated K_2P_-mediated leak currents and plasticity of IE in SST INs.

To test whether TRESK or other K2P channel genes are upregulated at the mRNA level, we assayed for transcript expression in SST INs using HCR-FISH, and found no increase in the expression of any of the Kcnk genes tested, tested at multiple time points after deprivation onset (Supp. Fig. 3). Thus, the TRESK channel is upregulated at the protein level but not the transcript level.

## Discussion

SST INs play a critical role in cortical circuits by providing dendritic inhibition to PYR cells that shapes sensory processing^8,55^ and regulates plasticity of PYR excitatory synapses. As a result, SST INs gate motor learning in motor cortex^56–58^, and sensory plasticity in visual cortex and S1^4,10^. In contrast, much less is known about experience-dependent plasticity in SST INs themselves. In V1, visual experience also regulates excitatory synaptic input to SST INs^13^, while in motor cortex, motor learning reduces SST inhibitory output synapses which is thought to enable motor learning in PYR networks^11^. Hippocampal O-LM interneurons exhibit altered IE in response to network stimulation during LTP and LTD induction protocols^17,18^. In S1, tactile stimulus-reward association learning weakens excitatory synaptic input to SST INs, which reduces SST activity to disinhibit PYR cells^14,15^. In L4 SST INs of S1, low-threshold spiking (LTS) SST INs have been shown to undergo differential plasticity of IE following classical sensory conditioning or pseudo-conditioning tasks^59^. While this illustrates plasticity of IE as a crucial mechanism of plasticity that can be engaged by learning, these experiments are limited to a subpopulation of SST INs. In addition, it is unclear whether naturalistic sensory experience drives plasticity of IE in SST INs.

We found that cortical L2/3 SST INs in S1 exhibit robust experience-dependent plasticity of intrinsic excitability following whisker deprivation. This effect, which was seen at 3 days of whisker deprivation, was apparent as a reduction in F-I curves due to reduced input resistance, without change in V_m_, spike threshold, or spike shape (Fig. 2). Comparing this effect to the normal development of SST IE revealed that this represents a *de novo* weakening as opposed to a developmental delay (Fig. 2C). Deprivation also slightly increased excitatory synaptic input to SST INs, evident as an increase in mEPSC frequency. To identify the biophysical mechanisms underlying the weakening of SST IE, we tested for changes in currents that had been previously associated with plasticity of IE in other cell types (K_V_1, HCN, K_V_7). None of these were altered by deprivation. Instead, deprivation caused an increase in outward leak K^+^ current, mediated by K_2P_ channels, which are a major contributor to resting K^+^ conductance. Thus, these findings identify K_2P_ channels as a novel locus for plasticity of IE, and the main site by which whisker deprivation modulates IE in SST INs.

The biophysical mechanisms for plasticity of IE that we observed in SST cells are distinct from those identified in other cell types. In PV INs in S1, whisker deprivation also reduces IE, but with differences in time scale and mechanism^16^. PV IE is reduced in deprived S1 columns after just 24 hours of whisker deprivation, in contrast to SST INs, which show reduced IE after 3 days of deprivation but not 1-2 days (Fig. 2 and Suppl. Fig. 1). Deprivation reduces F-I curves in PV INs by depolarization of spike threshold mediated by upregulation of the voltage-gated K^+^ channel, K_V_1.1, without any change in input resistance^16^. In contrast, SST INs show stable spike threshold and K_V_1.1 expression, and instead F-I curves are reduced by a decrease in input resistance that is mediated, at least in part, by upregulation of K_2P_ conductance. Thus, cortical INs display cell-type specific IE plasticity mechanisms in response to the same sensory manipulation. Mechanisms for IE plasticity in PYR cells are varied, but in S1 involve increased AIS length as a homeostatic response to increase PYR excitability in response to prolonged sensory deprivation^30^. In SST INs, AIS length was unaltered (Suppl. Fig. 2). This again indicates distinct, cell type specific mechanisms for IE plasticity.

This mechanism of IE plasticity we observed in SST INs in S1 is also different from O-LM cells in hippocampus, which are a well-studied SST IN subtype. Acute theta burst stimulation of CA1 PYR afferents to O-LM cells causes long-term potentiation of IE (LTP-IE) in O-LM cells^18^. This LTP-IE occurs due to a phospholipase C (PLC) dependent depletion of PIP_2_, ultimately leading to the downregulation of both K_V_7 and HCN currents^18^. Conversely, a negative spike pairing protocol drives long-term depression of IE (LTD-IE) in O-LM cells, due to elevated endocannabinoid signaling (eCB) causing an upregulation of K_V_7 current^17^. Deprivation-induced plasticity in S1 SST INs did not involve detectable modulation of K_V_7 or HCN currents (Fig. 5), so is a distinct mechanism. SST INs therefore seem to collectively express multiple biophysical mechanisms for IE plasticity, with different SST IN subtypes exhibiting distinct mechanisms, perhaps also dependent on induction protocol.

In a prior study on GIN neurons from neonatal-derived S1 cortical slice cultures, Gibson and colleagues found that chronic (2.5 day) activity blockade, by the application of TTX, leads to an increase in IE of GIN neurons^19^. This was evident as a shift in F-I curves accompanied by an increase in input resistance, which was due to downregulation of I_h_ and I_K(Leak)_^19^. Using pharmacology, they suggested the reduction in K_leak_ was due to reduction in K_2P_ channel conductance^19^, however GIN neurons comprise a subset of Martinotti-like SST cells (MCs)^33,35^. This may explain the general similarity in IE mechanisms, although IE plasticity in their study was opposite in sign to deprivation-induced plasticity at P17-19 in our study, which could reflect culture conditions or a layer- or age-dependent effect. Our finding that K_2P_ currents undergo plasticity in SST INs in response to whisker deprivation indicates that it is a physiologically relevant mechanism *in vivo*, likely recruited during natural shedding and regrowth of whiskers. Thus, our findings add to the breadth of plasticity mechanisms known to contribute to experience-dependent plasticity in cortex.

While we found altered expression of TRESK channel protein in SST cells after deprivation, a full molecular understanding of the regulation of K_2P_ channel(s) during plasticity of SST IE is lacking. In Gibson et al.’s slice culture experiments with TTX, they suggest based on pharmacology and pH sensitivity that TTX treatment regulates a K_2P_ channel in the TASK family^19^. Using publicly available single cell RNA-seq and Patch-Seq datasets^50,51^, we found that SST INs in cortex are enriched in TREK-1 (*Kcnk2*), TASK-1 (*Kcnk3*), and THIK-2 (*Kcnk12*), consistently across multiple datasets. While these datasets did not detect meaningful TRESK (*Kcnk18*) expression, we found using a commercially available antibody that TRESK expression occurs in L2/3 SST INs and is significantly upregulated after whisker deprivation (Fig. 6). The scRNAseq and PatchSeq datasets may have missed TRESK expression due to methodological issues or because TRESK may be expressed at low levels during normal sensory experience, and is only upregulated with sensory deprivation, which was not part of these earlier studies. Future studies are needed to understand mechanisms and prevalence of activity-dependent regulation of TRESK and other K_2P_ channels.

The net effects of reduced IE in SST INs on S1 cortical network function remain unknown. Because many L2/3 SST INs are involved in recurrent (feedback) inhibition of PYR cells, reduced SST IE may function as a slow homeostatic mechanism to restore mean firing rate of PYR cells, by reducing recurrent inhibition on PYR cells. However, this homeostatic regulation is slower than that implemented by PV cells^16,60^. Alternatively, the reduction in SST IE may primarily serve to locally disinhibit PYR dendrites, thus promoting LTP or changing plasticity rules at PYR-PYR excitatory synapses^58^. This could open a period of functional reorganization in L2/3 networks after several days of whisker deprivation, consistent with the onset of plasticity in whisker-evoked spiking in L2/3 after >3 days of D-row whisker deprivation^32^. In primary visual cortex, it has been reported that repeated exposure to varied visual stimuli elevates SST IN activity and enhances beta activity^12^. This occurs through a VIP-SST disinhibitory circuit, where repeated exposure to varied visual stimuli reduces VIP IN activity and subsequently increases SST IN activity. The net effect of this increase in dendritic inhibition is enhanced visual selectivity of PYR cells. Our results indicate that sensory experience also directly alters SST intrinsic function, which would alter SST output onto PYR cells, and thus may robustly shape PYR cell responsiveness to sensory input in S1 and other cortical areas.

## Supplemental figures

**Supplemental Figure 1.**
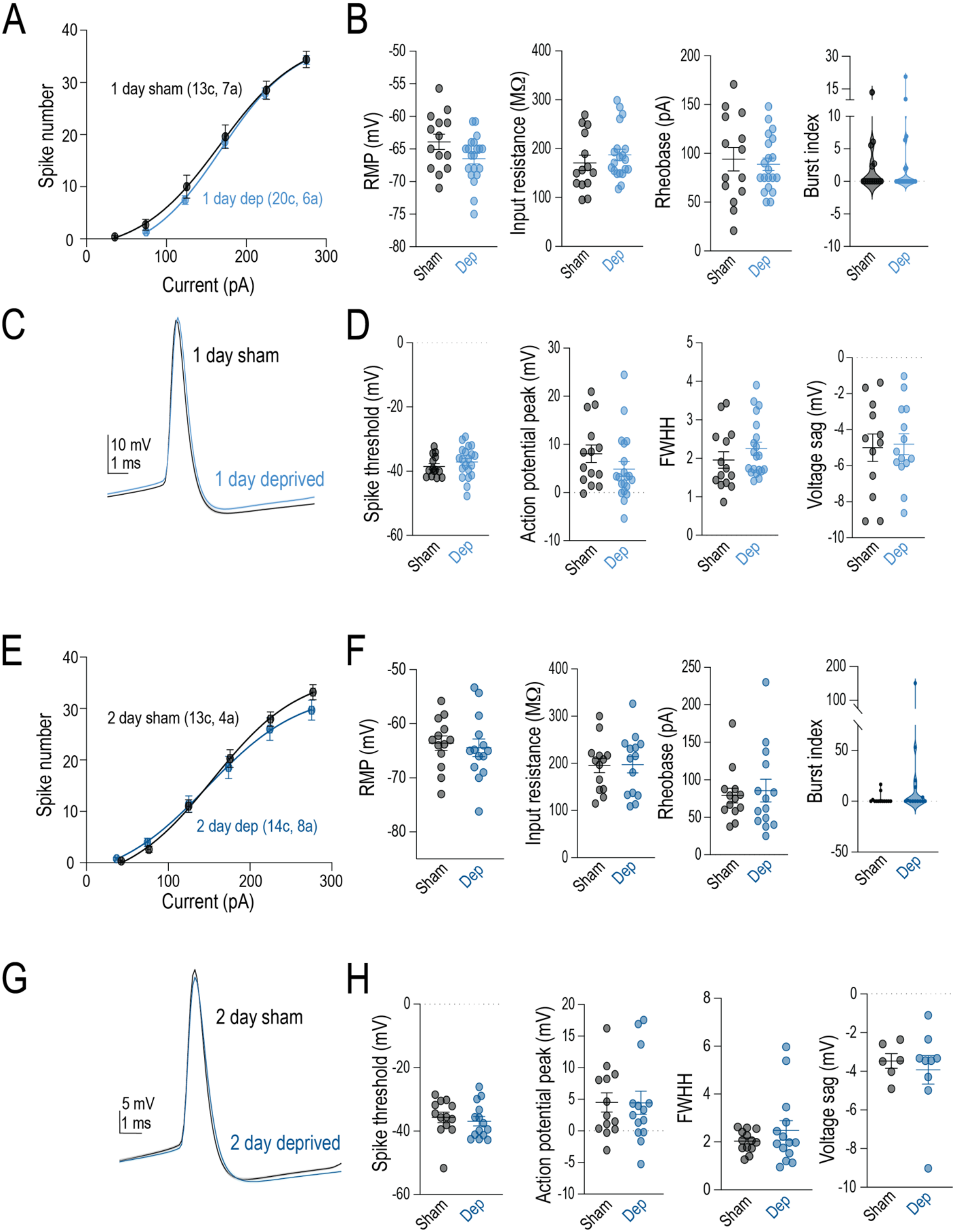
1- and 2-day whisker deprivation does not affect SST IE. **A.** Spike frequency-current (F-I) relationship comparing sham (black) and 1-day D-row deprived (light blue) mice. F-I curves were compared using 2-way ANOVA, p = 0.17, ns. **B.** Comparison of cell intrinsic properties including resting membrane potential (V_rest_), input resistance, and rheobase, as well as burst index in sham vs 1-day D-row deprived mice (Mann-Whitney rank comparisons, V_rest_ p = 0.15, ns; input resistance p = 0.29, ns; rheobase p = 0.82, ns; burst index p = 0.83, ns). **C.** Average spike waveform in sham vs 1-day D-row deprived mice. **D.** Comparison of spiking and active membrane properties including spike threshold, action potential peak, full width at half-height (FWHH), and voltage sag (Mann-Whitney rank comparisons, spike threshold p = 0.22, ns; action potential peak p = 0.14, ns; FWHH p = 0.18, ns; voltage sag p = 0.99, ns). **E.** Same as A, but for 2-day D-row deprivation. F-I curves were compared using 2-way ANOVA, p = 0.34, ns. **F.** Same as B, but for 2-day D-row deprivation (Mann-Whitney rank comparisons, V_rest_ p = 0.44, ns; input resistance p = 0.87, ns; rheobase p = 0.80, ns; burst index p = 0.24, ns). **G.** Same as C, but for 2-day D-row deprivation. **H.** Same as D, but for 2-day D-row deprivation (Mann-Whitney rank comparisons, spike threshold p = 0.30, ns; action potential peak p = 0.94, ns; FWHH p = 0.99, ns; voltage sag p = 0.99, ns).

**Supplemental Fig. 2.**
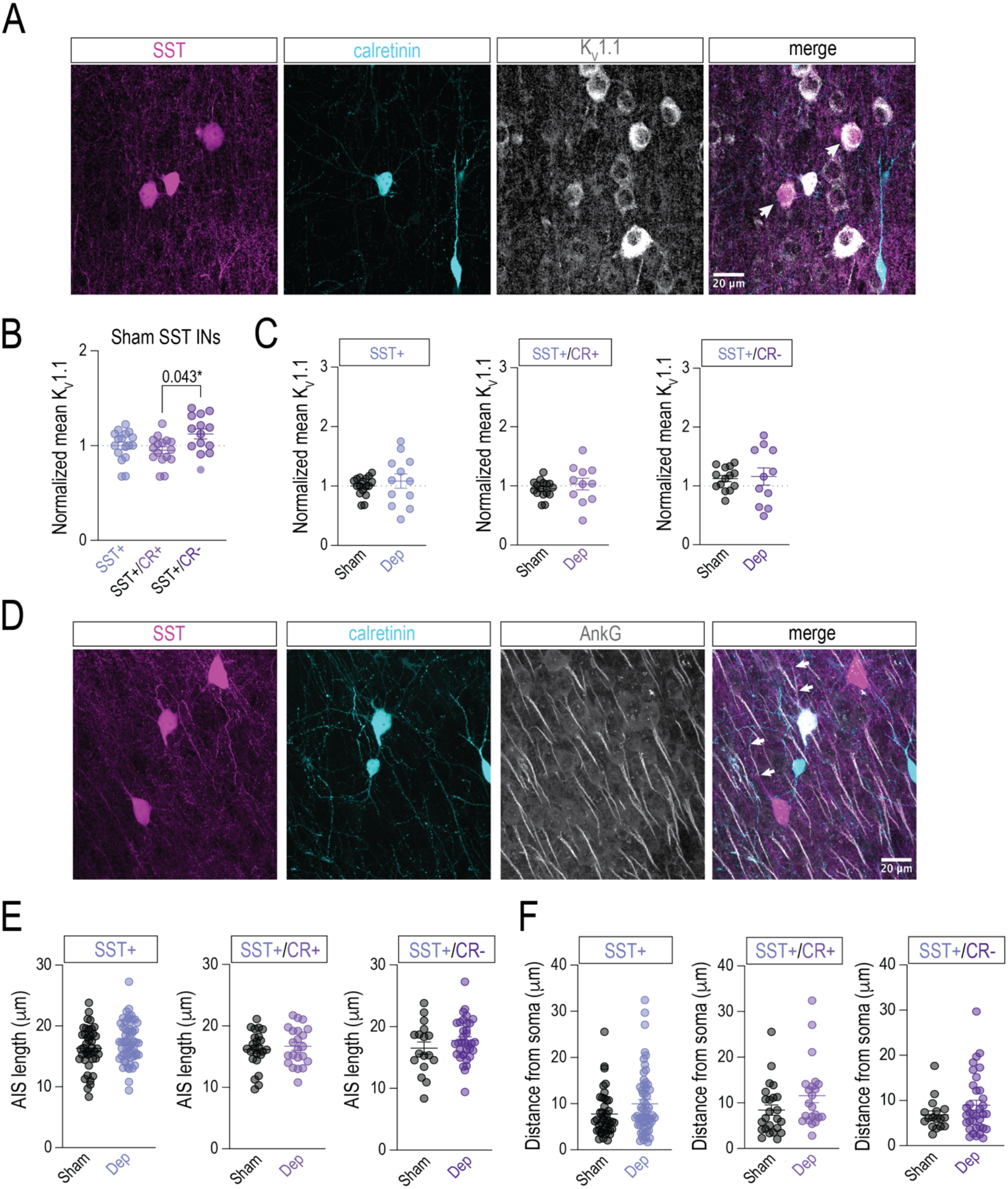
K_V_1.1 expression and axon initial segment (AIS) morphology are unaffected by whisker deprivation. **A.** Immunohistochemical co-staining of somatostatin (SST, magenta), calretinin (CR, cyan), and K_V_1.1 (gray). **B.** Mean fluorescence intensity of K_V_1.1 across subgroups of SST INs in sham mice (SST+: 1.00 ± 0.04, SST+/CR+: 0.951 ± 0.04, SST+/CR-: 1.123 ± 0.05; Kruskal-Wallis test: p =0.049*). **C.** Comparison of mean fluorescence intensity between SST IN subgroups in sham vs deprived mice (SST+ sham: 0.98 ± 0.03 vs SST+ deprived: 1.06 ± 0.12, Mann-Whitney rank comparison, p = 0.62, ns; SST+/CR+ sham: 0.92 ± 0.03 vs SST+/CR+ deprived: 0.97 ± 0.12, Mann-Whitney rank comparison, p = 0.40, ns; SST+/CR-sham: 1.09 ± 0.04, SST+/CR-deprived: 1.08 ± 0.14 Mann-Whitney rank comparison, p = 0.99, ns). **D.** Immunohistochemical co-staining of somatostatin (SST, magenta), calretinin (CR, cyan), and ankyrin G (AnkG, gray). State whether D is sham or deprived mouse. **E.** Comparison of axon initial segment (AIS) length (measured using AnkG staining) between SST IN subgroups in sham vs deprived mice (SST+ sham: 16.32 ± 0.54 μm vs SST+ deprived: 17.34 ± 0.43 μm, p = 0.14, ns; SST+/CR+ sham: 16.17 ± 0.62 μm vs SST+/CR+ deprived: 16.68 ± 0.65 μm, p = 0.57, ns; SST+/CR-sham: 16.52 ± 1.0 μm vs SST+/CR-deprived: 17.76 ± 0.58 μm, p = 0.26, ns). **F.** Comparison of the distance of the AIS from the soma between SST IN subgroups in sham vs deprived mice (SST+ sham: 7.76 ± 0.76 μm vs SST+ deprived: 9.98 ± 0.91 μm, p = 0.10, ns; SST+/CR+ sham: 8.42 ± 1.15 μm vs SST+/CR+ deprived: 11.59 ± 1.56 μm, p = 0.06, ns; SST+/CR-sham: 6.84 ± 0.86 μm vs SST+/CR-deprived: 8.96 ± 1.09 μm, p = 0.49, ns).

**Supplemental Fig. 3.**
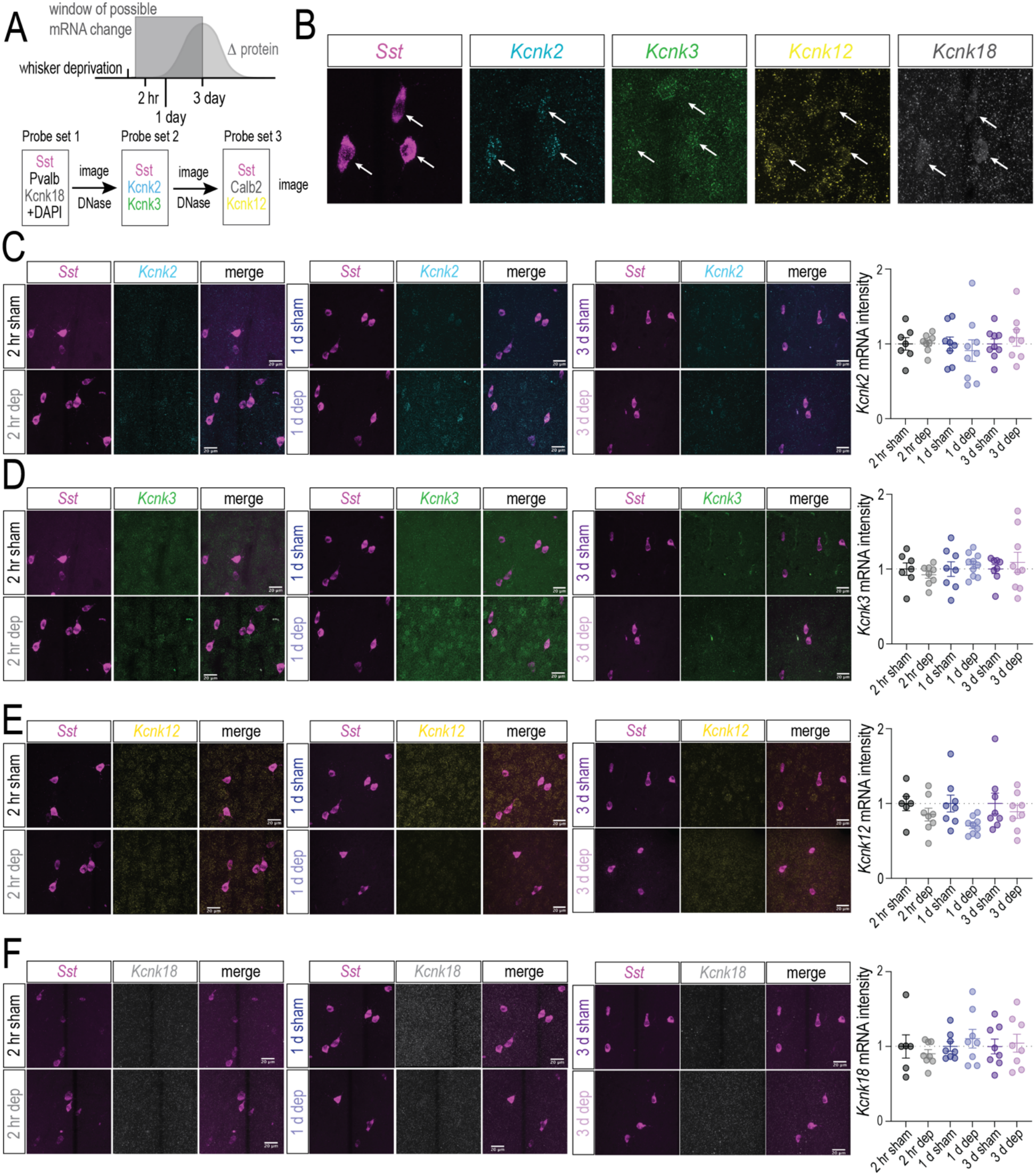
Unchanged expression of *Kcnk* gene transcripts following whisker deprivation. **A.** Schematic of hypothesized timing of mRNA vs protein changes across duration of whisker deprivation, and illustration of hybridization chain reaction fluorescence *in situ* hybridization (HCR-FISH) and imaging workflow. **B.** Representative images of different HCR-FISH labeling of *Sst* and *Kcnk* gene transcripts (*Sst, Kcnk2, Kcnk3, Kcnk12, and Kcnk18*). **C.** Representative images of *Kcnk2* gene expression after 2-hour, 1-day, and 3-day whisker deprivation (left) and quantification of fluorescence intensity (right). The fluorescence intensity of each *Kcnk* gene transcript was measured and then normalized to the average mean fluorescence intensity for each sham group (ex. the mean fluorescence intensity values of Kcnk2 for 2-hr sham and deprived were both normalized to the sham group. **D.** Same as C, but for *Kcnk3* gene expression. **E.** Same as C, but for *Kcnk12* gene expression. **F.** Same as C, but for *Kcnk18* gene expression.

## Acknowledgements

This work was supported by the National Institutes of Health (R01NS134639to D.E.F and F32 FNS119282A to B.M.B).

## Author contributions

BM. Bohannon was responsible for conceptualization of the project, execution of electrophysiological and molecular biological experiments, as well as writing and editing of the manuscript. K. Kaimaktchiev, C. Yang, NL. Alonso, M. Wang, and MG. Walker were responsible for performing immunofluorescence co-staining experiments, data analysis, and contributions to the writing of the methods section of the manuscript. DE Feldman provided project guidance and edited the manuscript.

